# WDL4 is a microtubule associated protein required for phytochrome B dependent thermomorphogenesis

**DOI:** 10.1101/2025.03.10.641267

**Authors:** Kristina Schaefer, Sidney L. Shaw

## Abstract

Flowering plants evolved an array of environmental sensors important for guiding adaptive morphological responses. The response to elevated ambient temperature involves the thermo-conversion of the light-sensing protein, PHYTOCHROME B (PhyB), leading to the activation of the nuclear transcription factor, PHYTOCHROME INTERACTING FACTOR 4 (PIF4). Here we employ temperature and light treatments to dissect the role of WAVE DAMPENED2-LIKE 4 (WDL4), a microtubule associated cytoplasmic protein, in modulating this signaling pathway. The WDL4 mutant (*wdl4-3*) phenocopies the loss of function *phyB-9* mutant at both 22° and 28° C for seedling and adult growth responses. Similarly, seedling hypocotyl elongation responses to red and far-red light exposure are strongly correlated between *phyB-9* and *wdl4-3*. Introduction of the *pif4-101* mutation into the *wdl4-3* background blocks *wdl4-3* hypocotyl hyper-elongation, indicating a specific PIF4 requirement. Addition of exogenous auxin, shown to rescue the *pif4* thermomorphogenetic response, restores hypocotyl elongation to wild type levels in the *pif4 wdl4-3* double mutant at 28° C, but fails to elicit the *wdl4-3* hyper-elongation phenotype indicating additional factors beyond auxin level modulating the thermomorphogenesis. Our data place a microtubule associated protein as a key regulator of PhyB-dependent thermomorphogenesis and photomorphogenesis response pathways.

## Introduction

Plants exposed to elevated ambient temperatures exhibit morphological and developmental changes that modulate effects on growth and life cycle, termed thermomorphogenesis (TMG) (Casal and Balasubramanian, 2019). Flowering plants above their normal temperature range typically exhibit hyper-elongation of the hypocotyl and petioles, hyponastic leaf elevation, and changes to bolting or flowering time (Casal and Balasubramanian, 2019). The latter results in changes to the timing of expressed developmental genes (Hatfield and Prueger, 2015) and the former is induced by changes in cell growth genes (Stavang et al., 2009). Genetic and molecular analyses of the TMG response in flowering plants have focused on the seedling hypocotyl owing to the easily visible hyper-elongation phenotype and the breadth of studies examining hypocotyl growth response to temperature, light, and other environmental cues (Krahmer and Fankhauser, 2024).

Evidence that elevated temperature elicits hypocotyl hyper-elongation through a genetic pathway was discovered in studies of the phytohormone auxin (Gray et al., 1998, Franklin et al., 2011). When grown at elevated temperature, the auxin co-receptor mutants, *axr2-1* and *axr1-12,* failed to show hyper-elongation, indicating a transcriptional requirement for elongation outside of the general changes in plant physiology related to temperature (Gray et al., 1998). More recent work has established phytochrome B (PhyB) as a key thermo-sensor controlling TMG (Legris et al., 2016, Jung et al., 2016). PhyB coordinates light and temperature signals to promote or repress cell elongation. PhyB is synthesized as a red-light absorbing form (Pr) that can be reversibly photoconverted to a far-red-light absorbing form (Pfr) to change the activity state (Casal and Balasubramanian, 2019). When red light is dominant, PhyB accumulates in the nucleus at foci linked to the inactivation of transcription factors in the Phytochrome Interacting Factor (PIF) family, known to influence hypocotyl elongation (Chen et al., 2003, Kim et al., 2023). When far-red light is elevated, PhyB no longer inactivates PIFs, resulting in hypocotyl elongation (Huq and Quail, 2002). Genetic and biochemical data showing that PhyB is specifically inactivated as a function of temperature (Jung et al., 2016, Legris et al., 2016, Burgie et al., 2021) established PhyB as an important control factor for TMG initiation (Jung et al., 2016, Legris et al., 2016, Burgie et al., 2021).

The PIF gene family member, *PIF4,* plays an essential role in TMG for the light-grown hypocotyl (Huq and Quail, 2002). Hypocotyl cells coordinately extend along a single axis, requiring both control of cell enlargement (i.e., growth) and regulation of cell shape (Gendreau et al., 1997, Krahmer and Fankhauser, 2024). PIF4, together with auxin-dependent, *ARF6* (Ulmasov et al., 1999), and brassinosteroid-dependent, *BRZ1* (Wang et al., 2012), appear to integrate the developmental and environmental stimuli impinging on the hypocotyl to direct elongation and trophic bending (Oh et al., 2014). At elevated temperatures, PhyB rapidly converts to the inactive Pr form (Jung et al., 2016, Legris et al., 2016) and PIF4 transcript levels become elevated (Koini et al., 2009). Whether TMG-induced morphological changes are solely dependent on PIF4 levels is unclear (Koini et al., 2009, Huq and Quail, 2002). PIF4, amongst other actions, directly regulates auxin biosynthesis genes (i.e. YUCCA 8, TAA1) (Franklin et al., 2011, Sun et al., 2012) together with auxin receptors and downstream signaling genes (TIR1/AFBs, AUX/IAAs, SAURs, etc.) (Pucciariello et al., 2018, Casal and Balasubramanian, 2019). Increased auxin production then promotes cell wall loosening via activation of proton pumps to induce wall acidification and cell enlargement (Fendrych et al., 2016, Spartz et al., 2012). Auxin has also been shown to induce cortical microtubule reorientation into patterns required for axial cell growth through the same auxin co-receptors required for TMG (True and Shaw, 2020, Elliott and Shaw, 2017). Hence, auxin has been proposed as a central regulator for TMG downstream of PhyB temperature sensing and PIF4 activation (Koini et al., 2009, Quint et al., 2016).

We previously characterized a member of the Arabidopsis *Wave Dampened2 Like* gene family (Yuen et al., 2003) and showed that WDL4 is a microtubule associated protein that negatively regulates hypocotyl elongation under lighted conditions (Schaefer et al., 2023). The phenotype was unusual in that a WDL4 mutant (due to tDNA insertion in the 3^rd^ intron, which will now be referred to as *wdl4-3*) failed to arrest hypocotyl growth between 4-5 days post-germination, leading to a hyper-elongation phenotype.

Moreover, the loss of WDL4 activity had no measurable effect on the cortical microtubule arrays in hypocotyl cells. Dark-grown seedlings on sucrose-free media had no significant growth phenotypes, but constitutive WDL4 expression retarded etiolated hypocotyl length by 30-40% (Schaefer et al., 2023). Interestingly, when *wdl4-3* seedlings were shifted from 22° C to 28° C under lighted conditions, we observed a dramatic hypocotyl hyper-elongation phenotype and elongated petioles. These observations led to the hypothesis that the microtubule associated WDL4 protein could be an important negative regulator of TMG in the hypocotyl. To address this hypothesis, we have evaluated *wdl4-3* growth under a variety of conditions and light perturbations using a loss-of-function PhyB mutant (*phyB-9*) as a comparator for TMG and phytochrome-dependent phenotypes.

## Results

### *wdl4-3* Seedlings Exhibit Thermomorphogenesis Phenotypes

To investigate the possible role of *wdl4-3* in TMG, we compared seedling growth phenotypes for wild type (Col-0), *wdl4-3*, a *wdl4-3* rescue expression line in which genomic WDL4 is expressed under its native promoter (WDL4_pro_:WDL4-mNEON *wdl4-3)*(Schaefer et al., 2023), and the loss of function *phyB-9* mutant (Reed et al., 1993). Seedlings were grown under continuous light at 22° C for 3 days before shifting to 28° C or remaining at 22° C. All plants were grown on sucrose free media due to the exaggerated wild type hyper-elongation at 28 ° C on sucrose supplemented media (**Supplemental Fig. 1**). Compared to wild type at 22° C, *phyB-9* seedlings displayed hyponastic leaves (measured as petiole angle), extended petioles, and shorter roots with exaggerated phenotypes observed at 28° C (**Supplemental Fig. 2**). Loss of function *wdl4-3* seedlings at 22° C exhibited extended petiole lengths, leaf hyponasty, and shortened root lengths comparable to *phyB-9.* Growth of *wdl4-3* at 28° C phenocopied the thermonastic responses observed for *phyB-9* (**Supplemental Fig. 2**). The WDL4_pro_:WDL4-mNeon returned hypocotyls to wild type length, with a small, significant residual difference in hypocotyl (0.92 +/- 0.1mm for WDL4_pro_:WDL4-mNeon vs 0.85 +/- 0.1 mm for wild type, p-value = 0.01) and petiole length at 22 °C compared to wild type (0.78 +/- 0.2 mm for WDL4_pro_:WDL4-mNeon vs 0.70 +/- 0.2 mm for wild type, p-value = 0.004). The WDL4_pro_:WDL4-mNeon line showed a similar response at 28° C when compared to wild type restoring nearly all of the wild type values (**Supplemental Fig. 2**). These data indicate that WDL4 negatively regulates the thermomorphogenic response in *Arabidopsis* seedlings.

### *wdl4-3* Plants Exhibit Thermomorphogenesis Phenotypes Throughout Adulthood

To determine the extent of WDL4 function in TMG, we examined TMG phenotypes for adult plants in wild type (Col0), *wdl4-3*, *phyB-9*, and WDL4_pro_:WDL4-mNeon *wdl4-3*. Seedlings grown on ½ MS plates were transferred to soil after 10 days and grown at 22° C or 28° C under 12-hour light and dark cycles through adulthood. Plants were imaged every 7 days (**Fig. 1A-D**). At 14 days, leaf hyponastic growth, controlled by asymmetrical cell elongation (Koini et al., 2009, van Zanten et al., 2009), was significantly higher in *phyB-9* and *wdl4-3* when compared to wild type and WDL4_pro_:WDL4-mNeon *wdl4-3* lines (**Fig. 1A-D**). Hyponasty appeared less evident by 3 weeks (**Fig. 1A-D**), likely due to the leaf weight and the weakness of the extended petioles. Wild type rosettes grown at 22° C for 21 days were compact, with petioles measuring 6.8 +/- 2.0 mm and a petiole angle at 72.8 +/- 9.0 degrees to the plant growth axis (**Fig. 1A, E, & F**). At 28° C, wild type petioles were stimulated to hyper-elongate to 12.8 +/- 2.9 mm (∼190%) and petioles were raised closer to the plant growth axis with angles at 58.1 +/- 14.3°, or a change of ∼15° (**Fig. 1A, E, & F**). At 22° C *phyB-9* and *wdl4-3* petioles were hyper-extended (13.5 +/- 5.2 and 15.4 +/- 5.8 mm, respectively) and hyponastic (59.0 +/- 20.1° and 59.3 +/- 19.2°) (**Fig. 1B, C, E, & F**). At 28° C, *phyB-9* and *wdl4-3* petioles were hyper-extended with respect to wild type, (13.9 +/- 3.7 mm and 17.3 +/- 4.9 mm respectively), but not significantly different from *phyb-9* or *wdl4-3* petioles at 22° C (**Fig. 1E**). Petiole angles at 28° C in *phyb-9* and *wdl4-3* were closer to the growth axis, decreased to 38.5 +/- 17.8° and 36.5 +/- 16.5°, respectively, a change of ∼20° and 23° (**Fig. 1F**). At 22° C, WDL4_pro_:WDL4-mNeon *wdl4-3* petiole length (7.8 +/- 2.2 mm) and petiole growth angle (71.3 +/- 13.1°) were not significantly different from wild type (**Fig. 1A, D, E, & F**). The WDL4_pro_:WDL4-mNeon *wdl4-3* line grown at 28° C had hyper-extended petioles to 10.83 +/- 3.59 mm, or an increased length of ∼140% and raised 57.01 +/- 15.02°, or a change of ∼14°. The growth phenotypes observed in all backgrounds were maintained through 6 weeks of growth. These data suggest WDL4 regulates petiole elongation and hyponasty in the adult plant response to elevated ambient temperatures.

**Figure 1:**
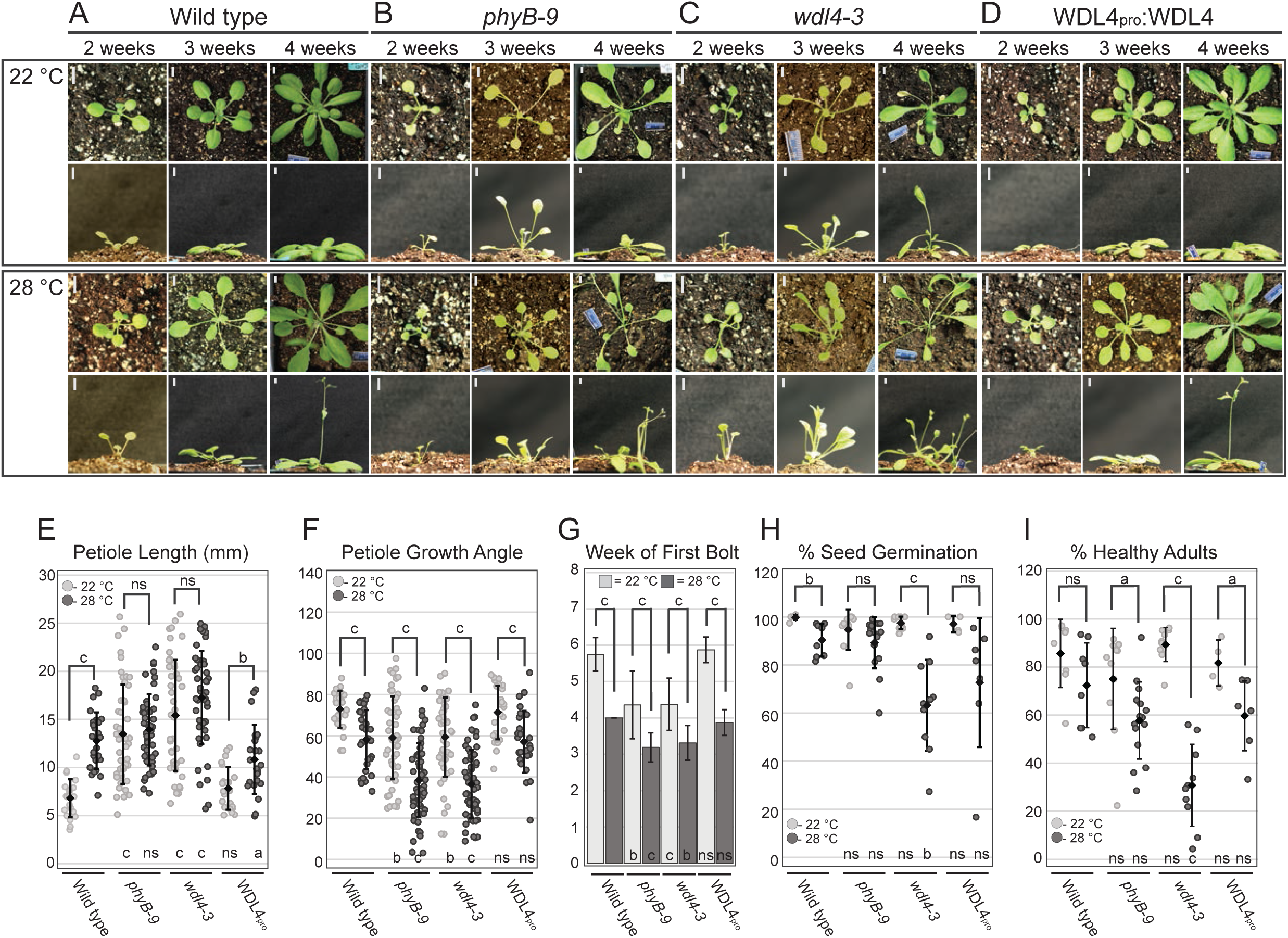
*wdl4-3* plants exhibit thermomorphogenetic phenotypes. Representative images of Arabidopsis Col-0 A) Wild type, B) *phyB-9,* C) *wdl4-3*, and D) genomic WDL4_pro_:WDL4-mNEON in the *wdl4-3* background (WDL4_pro_) grown in 12/12 hr light/dark for the indicated number of weeks and temperature. Scale bars = 5 mm. E) Petiole lengths measured at 3-weeks post-germi-nation at 28°C are significantly increased for wild type and rescue lines with no significant change in *phyB-9* and *wdl4-3* (n ≥ 21). F) Petiole hyponasty measured at 3 weeks post-germination increased (smaller angle) in all genotypes at 28°C compared to 22°C (n ≥ 25). G) Mean time to bolting (by week) at 28°C increased for all genotypes when compared to 22°C (n = 8-16 plants). H) Nearly all seed from plants grown at 22°C germinated (∼98%) while seed from plants grown at 28°C trended lower for seed germination (n ≥ 435). I) Seed from parent plants grown at 28°C that did germinate had lower probability of reaching maturity than plants derived from parents grown at 22°C (n ≥ 411). E-I) Two-tailed student t-tests were used to compare measurements between 22°C and 28°C within a genotype (brackets and letters above) and between wild type and mutant lines with the same treatment (letters below). P-values: a < 0.05, b < 0.005, c < 0.0005, ns > 0.05. In dot plots, ♦ = mean value. Scale bars indicate standard deviation of the mean.

### *wdl4-3* Plants Exhibit Developmental Thermomorphogenesis Phenotypes

Growth of *Arabidopsis* and other flowering plants at elevated ambient temperature accelerates bolt emergence time and seed development/production (Casal and Balasubramanian, 2019). At 22° C, wild type plants initially bolted between 5 and 6 weeks of germination. At 28° C, bolting initiated at 4 weeks (**Fig. 1A & G**). The TMG mutant, *phyB-9,* bolted at 4-5 weeks at 22° C and was pushed to an earlier bolting at 3-4 weeks at 28° C (**Fig. 1B & G**). The *wdl4-3* mutant bolted between 4-5 weeks at 22° C and 3-4 weeks at 22° C, similar to *phyb-*9 (**Fig. 1C & G**). Expression of the ectopic

WDL4_pro_:WDL4-mNEON transgene in *wdl4-3* rescued the early bolting phenotype (**Fig. 1D & G**). All lines responded to elevated temperatures by bolting at least a week earlier, suggesting that even though *wdl4-3 and phyB-9* at 22° C bolt as though they are at 28° C, the plants still receive and respond to the elevated temperature signal.

To test effects on seed viability, seed was collected from individual plants and grown on ½ MS plates for 7 days. The percent germination was calculated and each germinated seedling was then classed after 10 days as viable or failed based on the appearance of root elongation and emergence of true leaves. Seed collected from wild type plants at 22° C and 28° C showed a significant decrease in germination (99.8 +/- 1% vs. 90.3 +/- 7%, p-value = 0.002), but no significant change in seed viability (85.7 +/- 14% vs. 72.4

+/- 18%, p-value = 0.120) (**Fig. 1H & I**). Seed from *phyB-9* showed an insignificant decrease in germination percentage at 22° C and 28° C compared to wild type but showed a significant difference in percentage of viable seedlings at 28° C (**Fig. 1H & I**). Seed from *wdl4-3* plants grown at 22° C germinated at a similar percentage to wild type plants (97.1 +/- 3% vs 99.8 +/- 1% respectively), however seed from plants grown at 28° C had a significant decrease in both germination and viability (**Fig. 1H & I**). The WDL4_pro_:WDL4-mNeon *wdl4-3* line germination percentage was not significantly different from wild type at 22° C (97.0 +/- 4%, p-value = 0.055) or 28° C (72.7 +/- 27%, p = 0.095) and the transgene fully rescued the seed viability phenotype when compared to wild type (**Fig. 1H & I**). Taken together, these data suggest that WDL4 functions in a signaling pathway that slows development and seed production at elevated temperatures.

### Far-Red Light Inhibits *wdl4-3* Hyper-Elongation

Our comparison of *wdl4-3* growth and developmental phenotypes with a mutant for a central temperature sensor, PhyB (Jung et al., 2016, Legris et al., 2016), indicated a strong overlap in function between these genes. From these results, we hypothesized that WDL4 could be acting on TMG through a PhyB-dependent pathway as opposed to being an independent temperature sensor. PhyB is also a light sensor and *phyB-9* has been shown to have an aberrant response to light treatment. We hypothesized that if WDL4 acted through a PhyB-dependent pathway, then *wdl4-3* would have a similar aberrant response to light treatment (Reed et al., 1993, Franklin and Whitelam, 2005). To evaluate this hypothesis, we tested the *wdl4-3* seedling hypocotyl response to supplemental far-red light (i.e., shade avoidance response) when grown at normal (22° C) temperature. We treated light-grown wild type and *wdl4-3* seedlings with supplemental far-red light (sFR, 735 nm, see Methods) for 5 days at 22° C and then measured hypocotyl lengths. Wild type hypocotyls elongated ∼175% compared to seedlings grown in constant white light (WL), indicating a significant shade avoidance response (WL: 2.0 +/- 0.4 mm vs. sFR: 3.4 +/- 0.9 mm **Fig. 2A & C**). Control *phyB-9* hypocotyls are relatively elongated (5.8 +/- 1.5 mm) in WL and were slightly shorter when treated with sFR (5.4 +/- 1.4 mm, **Fig. 2A & C**). *wdl4-3* hypocotyls under WL hyper-elongated to 6.1 +/- 1.4 mm and were non-significantly changed in sFR, reaching 5.8 +/- 1.8 mm (**Fig. 2A & C**). The WDL4_pro_:WDL4-mNeon *wdl4-3* line was not significantly longer in cWL compared to wild type and elongation was induced to ∼180% with sFR, a similar response to wild type (WL 2.2 +/- 0.5 mm vs sFR 4.0 +/- 1.2 mm, **Fig. 2A & C**). Constitutive WDL4 expression (35s_pro_:WDL4-mNEON) had no significant effect on hypocotyl elongation in WL light, as previously determined (**Figure 2A & C**)(Schaefer et al 2023). However, sFR treatment of 35s_pro_:WDL4-mNeon seedlings resulted in an attenuated growth response ∼120% compared to wild type (**Figure 2A & C**, 1.8 +/- 0.6 mm and 2.2 +/- 1.2 mm). These data suggest that over-expression of WDL4 represses sFR induced hypocotyl hyper-elongation. These experiments support the hypothesis that WDL4 likely acts in a general PhyB pathway and is not acting specifically in a temperature response pathway.

**Figure 2:**
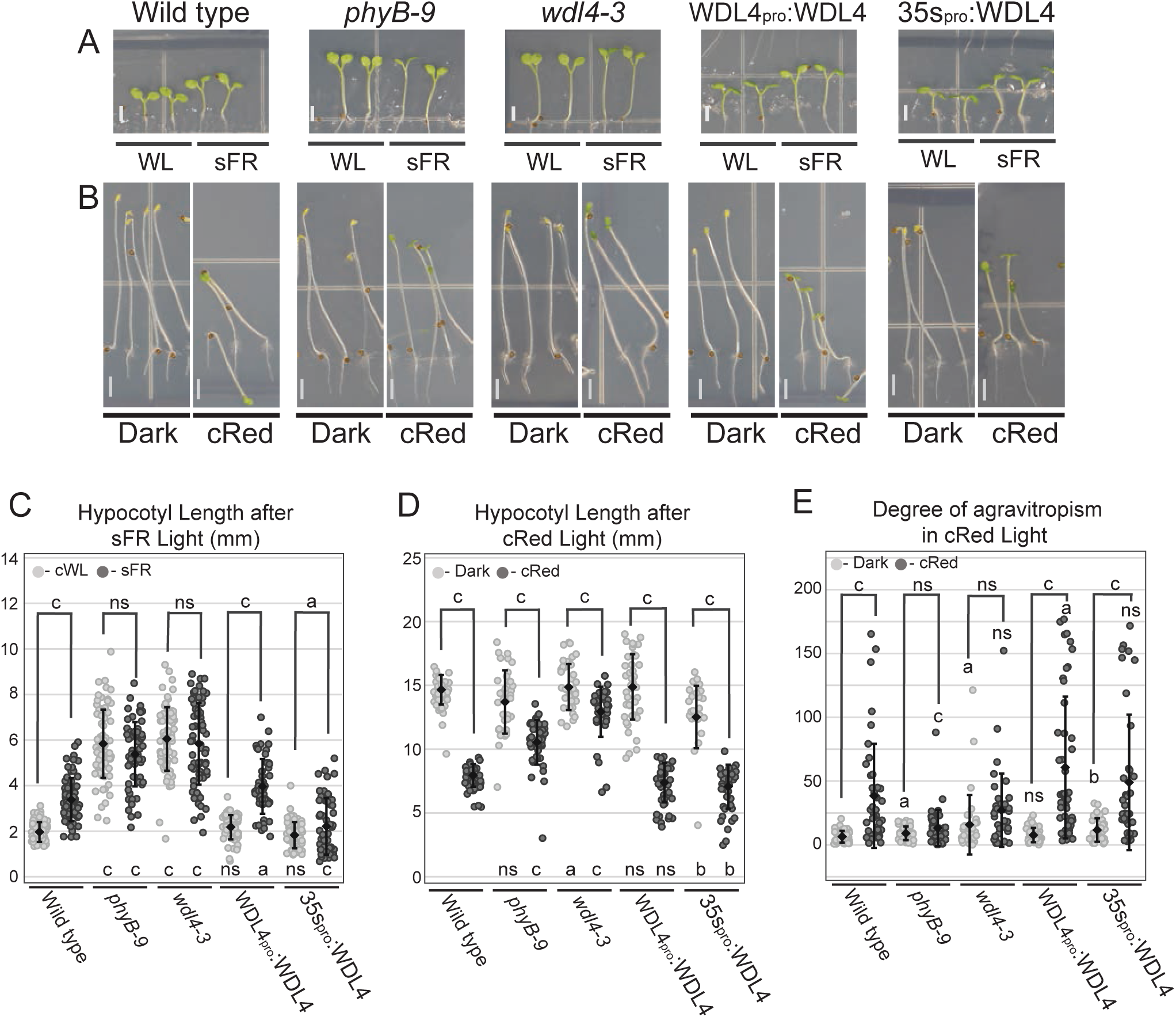
*wdl4-3* shows a decreased response to supplemental Far-red and red lights. Representative images of seedlings at A) 5 dpg in white light (WL, left) or WL supplemented with Far-Red light (sFR, ∼20 µmol m^-^² s^-1^ 735 nm, right). B) Seedlings at 4 dpg in darkness (left) or continuous red light (cRed, 20 µmol m^-^² s^-1^ 670 nm, right). Scale bar = 2 mm. C) sFR significantly increased hypocotyl length in wild type, WDL4_pro_:WDL4-mNEON *wdl4-3* (WDL4_pro_:WDL4) and 35s_pro_:WDL4-mNeon (35s_pro_:WDL4) lines but not *phyB* and *wdl4-3*. D) cRed significantly reduced hypocotyl elongation in all genotypes, but to a lesser degree in *phyB-9* and *wdl4-3* (reduction to 54% in wild type, 77% in *phyB-9*, 86% in *wdl4-3* of dark grown controls) (n ≥ 36). Changes in E) gravitropism (0° = vertical) were more pronounced in wild type, WDL4_pro_:WDL4-mNEON *wdl4-3* and 35s_pro_:WDL4-mNeon lines than in *phyB-9* and *wdl4-3*. C-E) Two-tailed student t-tests were used to compare measurements between control (i.e. WL or Dark) and light treatment within a genotype (brackets and letters above) and between wild type and the mutant line with the same treatment (letters below). P-values: a < 0.05, b < 0.005, c < 0.0005, ns > 0.05. In dot plots, ♦ = mean value. Scale bars indicate standard deviation of the mean.

### WDL4 Acts in Red-Light Induced Photomorphogenesis

Dark-grown etiolated seedlings can be triggered to undergo de-etiolation through red-light signaling of PhyB (Reed et al., 1994, Huq and Quail, 2002). To determine if WDL4 acts in this phytochrome-dependent signaling process, we grew wild type, *phyb-9*, *wdl4-3*, WDL4_pro_:WDL4-mNeon *wdl4-3*, and 35s_pro_:WDL4-mNEON seedlings in the dark or 670 nm red light (fluence – 20 µmol m^-2^ s^-1^) for 4 days. Wild type hypocotyls reached a length of 14.3 +/- 1.2 mm in darkness but only reached 7.7 +/- 0.9 mm in red light (**Fig. 2B & D**). Red light further induced cotyledon opening and color change from yellow to green (**Fig. 2B**). Control *phyB-9* hypocotyls grown in darkness were not significantly different from wild type (13.8 +/- 2.5 mm, student t-test p-value > 0.5) (**Fig. 2B**). Red light induced cotyledon opening and greening in *phyB-9* was comparable to the wild type response. Continuous red-light treatment reduced hypocotyl length in *phyb-9* (10.6 +/- 1.8 mm) when compared to dark grown, but growth retardation was significantly less than wild type, as previously reported (Lu et al., 2015). Dark-grown *wdl4-3* hypocotyls were slightly longer than wild type (15.0 +/- 1.8 mm, p-value = 0.03) and showed a subtle, but significant shortening, with far-red light (12.7 +/- 2.3 mm) (**Fig. 2B & D**). However, the growth attenuation in *wdl4-3* was significantly less than that observed in *phyb-9* (reduction of 86% and 77% respectively). Rescue line hypocotyls (WDL4_pro_:WDL4:mNeon *wdl4-3*) grew to 14.8 +/- 2.6 mm in darkness. Red light significantly retarded hypocotyl elongation comparable to wild type (7.2 +/- 1.6 mm, p-value > 0.5) (**Fig. 2B & D**). These data indicate that WDL4 plays a role in hypocotyl growth control in response to red-light signaling during de-etiolation.

### Red-Light Promotes Agravitropic Hypocotyl Elongation in *wdl4-3*

We observed that dark-grown *wdl4-3* hypocotyls showed aberrant gravitropism under red-light compared to wild type (**Fig. 2B**). Red light represses negative gravitropism and recent reports have suggested cytoplasmic PhyB regulates this response (Hughes, 2013, Hu and Lagarias, 2024). We hypothesized that *wdl4-3* seedlings are impaired for the red-light dependent repression of negative gravitropism. To test this hypothesis, we grew wild type and *wdl4-3* seedlings in darkness or in red-light using *phyB-9* as a control for red-light inactivation of negative gravitropism. Wild type, *phyB-9, wdl4-3,* WDL4_pro_:WDL4-mNeon *wdl4-3* and 35s_pro_:WDL4-mNeon grown in darkness showed roughly equivalent mean hypocotyl growth trajectories using the angle of the upper 1/3 of the seedling to evaluate negative gravitropism (range 5-15 degrees from 0) (**Fig. 2B & E**). Red-light treatment (20 µmol m^-2^ s^-1^) increased deviation from the mean growth vector in wild type an average of 38.5°, with a median of 24.0° (**Fig. 2E**). In contrast, gravitropism was not significantly disrupted by red light in *phyB-9* hypocotyls (mean 13.3 +/- 14,2°, median 11.3°, **Fig. 2E**) indicating an insensitivity to red light for this response. In the dark, *wdl4-3* growth trajectories had a slightly larger variance from 0° compared to *phyB-9* (mean = 15.9 +/- 23.3°) (**Fig. 2E**). Comparable to *phyB-9*, red light had no significant effect on gravitropism of the *wdl4-3* mutant (mean 27.2 +/- 28.7°, student t-test p = 0.1631). The WDL4_pro_:WDL4-mNeon *wdl4-3* line had similar growth trajectories in the dark (average 7.7 +/- 5.6°, median 5.7°) and an increased range in red light, compared to wild type (average 60.7 +/- 55.5°, median 38.2°) (**Fig. 2B & E**). Even though gravitropism was slightly but significantly altered in 35s_pro_:WDL4-mNeon dark grown seedlings (average 11.7 +/- 9.3°, median 9.7), red light growth trajectories were comparable to wild type (49.0 +/- 53.1°, median 24.4)( **Fig. 2B & E)**. These experiments suggested *wdl4-3* is not responding to red light signals related to negative gravitropism.

### *wdl4-3* Seedling and Rosette Growth Phenotypes Require PIF4

PhyB physically interacts with PIFs in the nucleus to inhibit their action on gene expression (Huq and Quail, 2002, Ni et al., 1999). Integration of the *phyB-9* allele into *pifq* (*pif1 pif3 pif4 pif5* quadruple mutant)(Leivar et al., 2012, Lee et al., 2021) background results in a short hypocotyl phenotype, indicating a requirement for PIF activity in the *phyB-9* hyper-elongation response (Huq and Quail, 2002). Multiple experiments indicate that elevated temperatures release PhyB repression of PIF4 to promote hypocotyl elongation (Stavang et al., 2009, Koini et al., 2009). Interestingly, double *phyB pif4* mutants exhibit elongated hypocotyls in the light (Huq and Quail, 2002, de Lucas et al., 2008). We hypothesized from our results that WDL4 could play a specific role in the PhyB/PIF4 pathway by negatively regulating the action of PIF4 on hypocotyl growth. Therein, we asked if the loss of PIF4 would block the hypocotyl hyper-elongation observed in the *wdl4-3* background at 22° and 28° C. We crossed in the Col-0 loss-of function PIF4 mutant, *pif4-101* (*pif4,* (Lorrain et al., 2008)), to *wdl4-3*, to create the *pif4 wdl4-3* double mutant.

Loss of function *pif4* seedlings grown at 22°C in the light had comparable petiole growth angles and root lengths to wild type but displayed shorter petioles (**Supplemental Fig. 3**). At 28°C, temperature-induced TMG phenotypes were reduced in *pif4* (**Supplemental Fig. 3**). At 22°C and 28°C *pif4 wdl4-3* double mutant seedlings exhibited the *pif4* single mutant phenotypes (**Supplemental Fig. 3**). These data indicate that *wdl4-3* seedlings require PIF4 to affect the hypocotyl hyper-elongation and related TMG phenotypes in the *Arabidopsis* seedling.

Adult *pif4* plants grown at 22°C on a 12/12 hr light regime were indistinguishable from their wild type counterparts (**Fig. 3A-B**). At 28° C, *pif4* petioles had a significantly reduced thermonastic response (**Fig. 3B & E**). At 22°C, the petiole hyper-elongation and hyponasty observed in *wdl4-3* was repressed in the double *pif4 wdl4-3* mutant (**Fig. 3C, E, & F and Fig. 1**). At 28° C, *pif4* plants bolted earlier than at 22° C and had an insignificant reduction in the number of seedlings surviving to adulthood, similar to wild type (**Fig. 3B, E, & I**). At 28° C, *pif4 wdl4-3* plants were more similar to *pif4*, in that there was little change in petiole elongation and hyponasty. These data suggest that PIF4 regulates petiole length at elevated temperatures and plays a minimal or over-lapping role in production of first bolt and seed fecundity in our experimental conditions. At 28° C, the double mutant line had a significant reduction in seed fecundity that was not seen in wild type or the *pif4* single mutant, resembling the effect observed in *wdl4-3* (**Fig. 3I vs Fig. 1I**), suggesting the reduction seed fecundity in *wdl4-3* is PIF4 independent.

**Figure 3:**
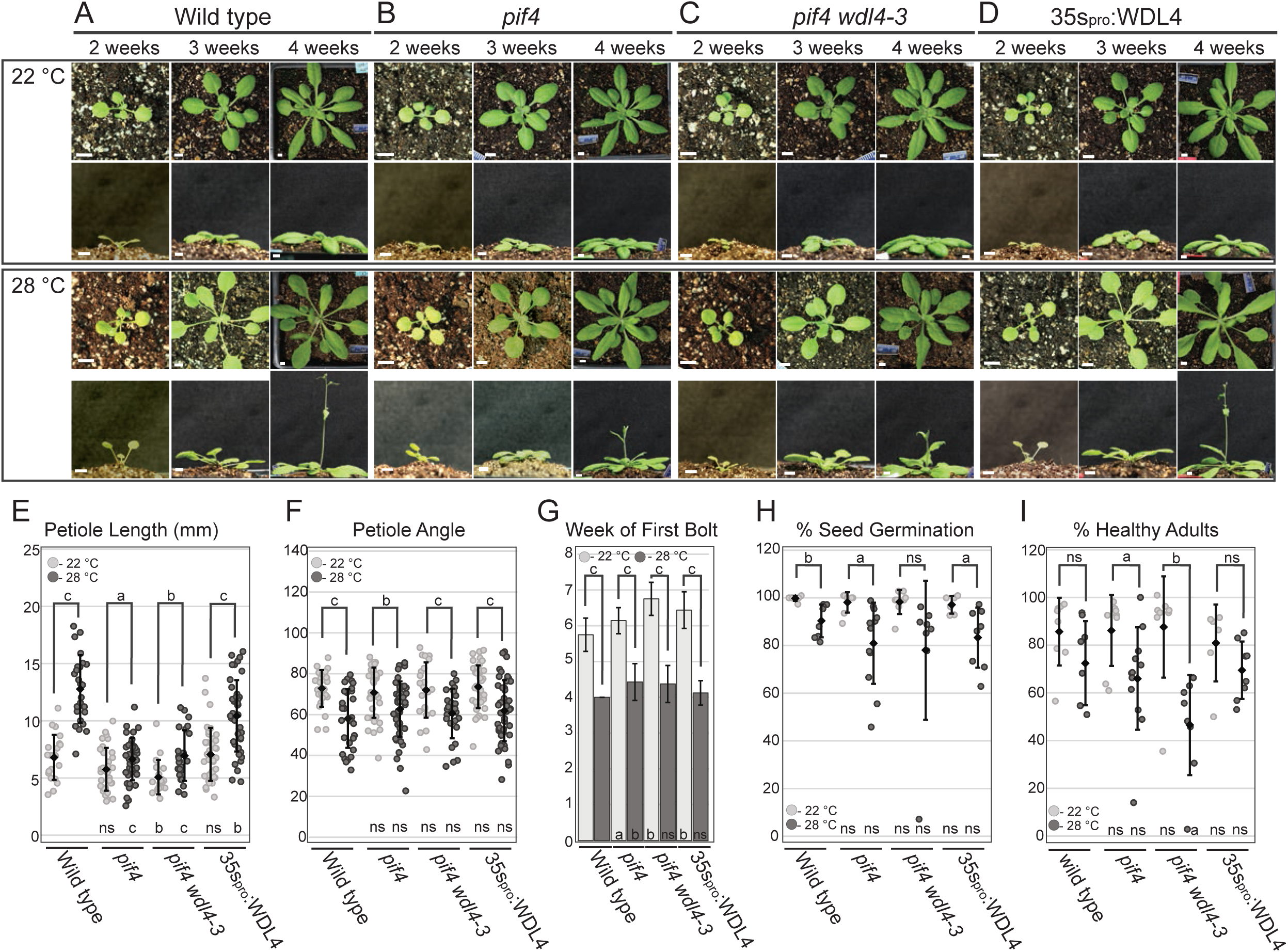
Loss of PIF4 eliminates *wdl4-3* adult thermomorphogenetic phenotype. Representative images of plant growth for A) wild type, B) *pif4*, C) *pif 4 wdl4-3*, and D) constitutive expression of genomic WDL4-mNeon via a 35sCaMV promoter (35s_pro_:WDL4). Plants were grown in 12/12 hr light/dark cycle and imaged on the indicated day. (Note: these experiments were performed together with experiments in Figure 1 and are compared to the same wild type datasets (E-I)). Scale bars = 5 mm. E) Petiole elongation was reduced in the *pif4* mutant when compared to wild type (n ≥ 19). F) Growth at 28°C increased petiole hyponasty (0° = direction of growth) to a similar extent in all genotypes (n ≥ 23). G) Growth at 28°C induced earlier bolting in all genotypes (n = 8-16 individual plants). H) Seed from parents grown at 22°C appeared unaffected for germination but depressed when parent was grown at 28°C (n ≥ 655). I) The percent of germinated seed producing seedlings with true leaves was not significantly altered in these genotypes when parent was grown at 28°C (n ≥ 635). E-I) Two-tailed student t-tests were used to compare measurements between 22°C and 28°C within a genotype (brackets and letters above) and between wild type and the mutant line with the same treatment (letters below). P-values: a < 0.05, b < 0.005, c < 0.0005, ns > 0.05. In dot plots, ♦ = mean value. Scale bars indicate standard deviation of the mean.

### WDL4 Red and Far-Red Light Phenotypes require PIF4

Supplemental Far-Red light (i.e., shade avoidance) induces hypocotyl elongation in light grown seedlings through PhyB and multiple PIF family members, including PIF4 (Lorrain et al 2008). Consistent with this work, we found that sFR signaled wild type hypocotyls to elongate to ∼200% of untreated at 22° C, where *pif4* hypocotyls were induced to grow, but only to ∼170% of untreated (*pif4* 1.1 +/- 0.2 mm in WL, 1.8 +/- 0.4 mm in sFR) (**Fig. 4A & C**). To determine if WDL4 altered this PIF4-dependence in the shade-avoidance response, we assayed the *pif4 wdl4-3* double mutant and observed a ∼130% increase in hypocotyl length after sFR treatment at 22° C (WL: 1.1 +/- 0.2 mm; sFR: 1.5 +/- 0.3 mm) (**Figure 4A & C**). Hence, the absence of WDL4 activity significantly increased the impact of PIF4 loss on far-red light dependent hypocotyl elongation (Student’s T-test, p-value < 0.0005).

**Figure 4:**
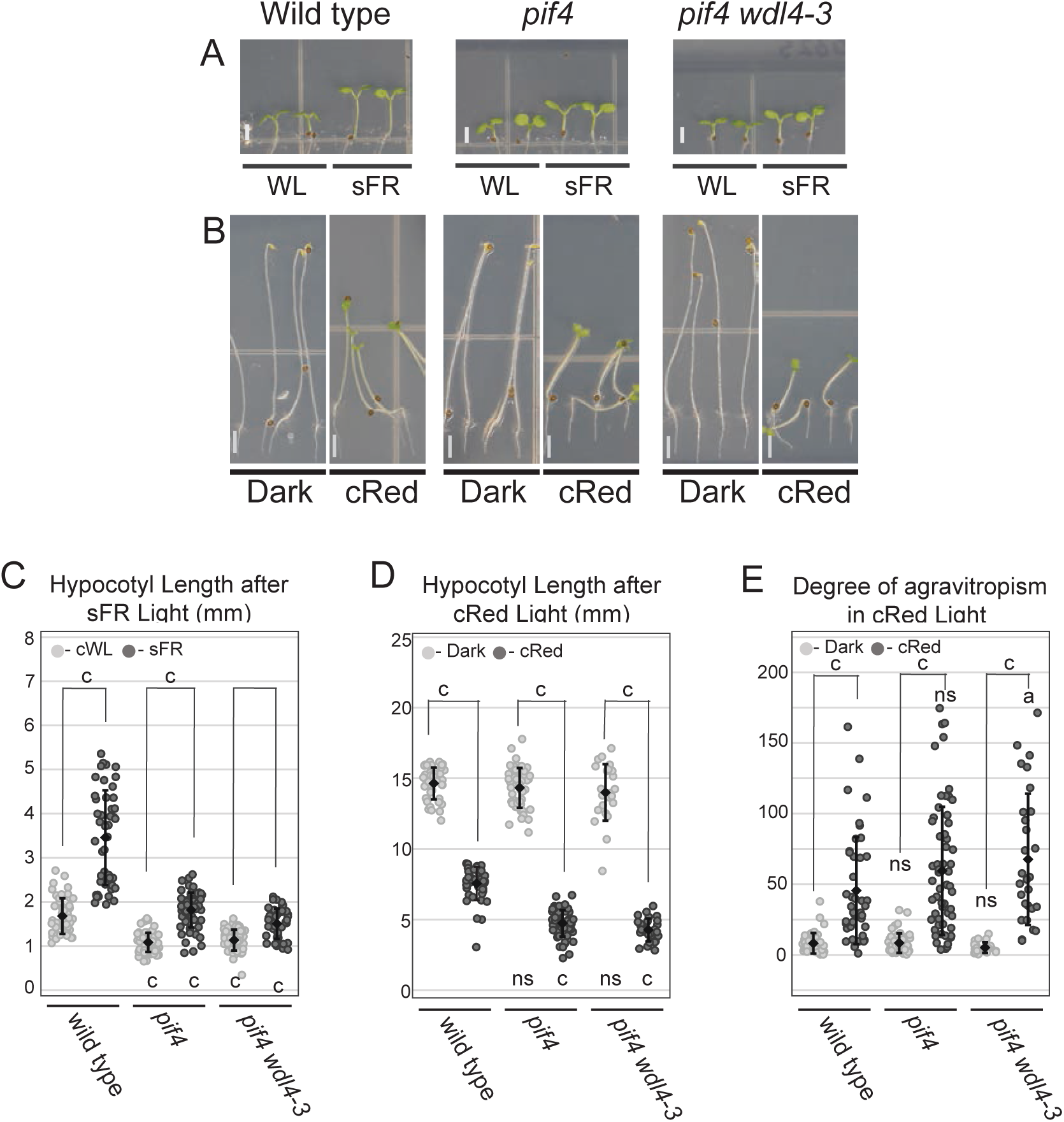
Loss of PIF4 restores *wdl4-3* response to sFR and Red light. Representative images of seedlings A) at 5 dpg in white light (WL - left) or WL supplemented with Far-red light (sFR, ∼20 µmol m^-2^ s^-1^ 735 nm, right). B) Seedlings at 4 dpg in darkness (left) or continuous red light (cRed, 20 µmol m^-2^ s^-1^ 670 nm, right). Scale bar = 2 mm. C) Hypocotyl length after sFR was significantly increased in all genotypes, but to a lesser extent in *pif4* and *pif4 wdl4-3* backgrounds (n ≥ 41). D) cRed significantly reduced hypocotyl elongation in all genotypes (n ≥ 24). E) Red light induced changes in gravitropism (0° = vertical) was more similar to wild type in *pif4 wdl4-3*, but not fully restored. C-E) Two-tailed student t-tests were used to compare measurements between control (i.e. WL or Dark) and light treatment within a genotype (brackets and letters above) and between wild type and the mutant line with the same treatment (letters below). P-values: a < 0.05, b < 0.005, c < 0.0005, ns > 0.05. In dot plots, ♦ = mean value. Scale bars are standard deviation of the mean.

Photomorphogenesis is induced in dark-grown wild type seedlings by cR treatment, reducing hypocotyl elongation by ∼50% (**Figure 4B & D**). In agreement with prior findings (Leivar et al., 2008), *pif4* hypocotyl elongation was more strongly reduced with cR light treatment, producing seedlings that were 33% of the dark grown counterparts (4.7 +/- 0.9 mm, **Figure 4D**). Double mutant *pif4 wdl4-3* seedlings phenocopied *pif4* hypocotyl lengths in both the dark and with cR treatment, reaching 30% of dark grown controls (Dark: 14.0 +/- 2.0 mm vs cR: 4.3 +/- 0.8 mm; **Figure 4B &D**). The cR treatment produced no significant difference in gravitropism response for the *pif4* mutant compared to wild type (**Figure 4E** Student t-test p-value = 0.077). The *pif4 wdl4-3* double mutant showed, however, a slight but significant agravitropic response to cR light when compared to wild type (**Figure 4E**, p-value = 0.035). These data indicate that the *wdl4-3* phenotypic responses to red light are PIF4 dependent.

### Auxin rescues *pif4* growth at 28° C, but does not recover *wdl4-3* hyper-elongation

PIF4 promotes transcription of auxin biosynthesis genes at elevated temperatures (Sun et al., 2012), and is proposed to increase auxin levels to induce cell elongation (Gray et al., 1998, Spartz et al., 2012). In support of this hypothesis, the shortened *pif4* hypocotyl elongation observed at 28° C can be restored to wild type response with exogenous auxin (Franklin et al., 2011). These experiments led to the hypothesis that TMG is driven by auxin upregulation, downstream of PIF4 (Franklin et al., 2011, Gray et al., 1998) Since PIF4 is clearly required for observed hyper-elongation in *wdl4-*3, we asked if exogenous auxin would rescue the 28° C light-grown *pif4 wdl4-3* hypocotyl back to wild type length, to the much longer *wdl4-3* length, or fail to rescue. To address this question, we grew Col-0, *phyB-*9, *wdl4-*3, *pif4*, and *pif4 wdl4-3 s*eedlings at 22° C for 3 days and then transferred seedlings to picloram plates (PIC – a thermostable auxin analog) at either 22 °C or 28 °C (**Fig. 5A**).

**Figure 5:**
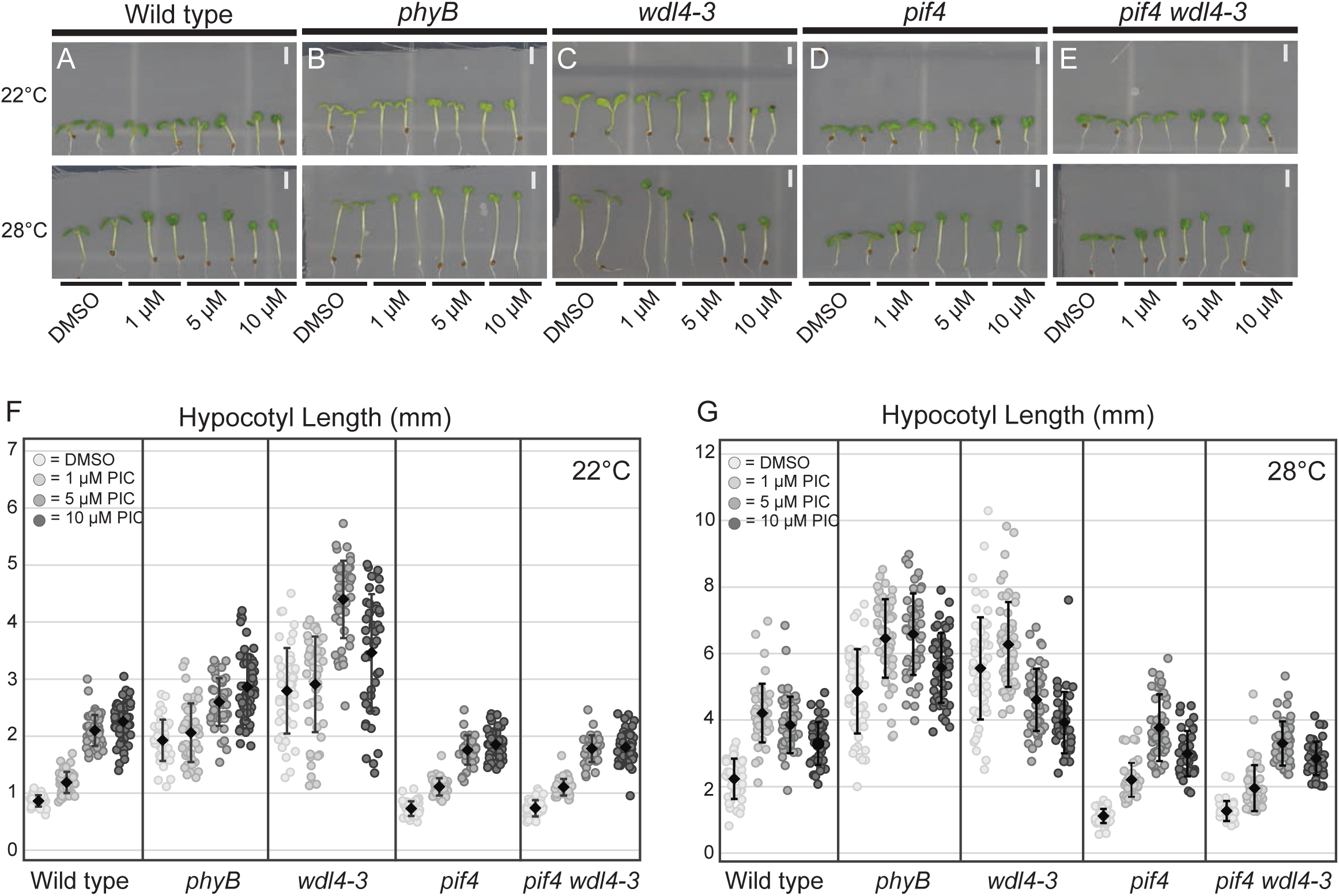
Supplemental auxin restores *pif4 wdl4-3* elongation to wild type levels at 28°C. Representative images of seedlings A) wild type, B) *phyB*, C) *wdl4-3*, D) *pif4*, and E) *pif4 wdl4-3* seedlings grown for 3 days on ½ MS media before being shifted to plates supplemented with DMSO, 1 µmol, 5 µmol, or 10 µmol of picloram and grown for an additional 3 days at 22 °C (top) or moved to 28 °C (bottom). Scale bar = 2 mm. Quantification of hypocotyl lengths after picloram treatment at F) 22 °C or G) 28 °C. Hypocotyls measured ≥ 37. In dot plots, ♦ = mean value. Scale bars are standard deviation of the mean.

Consistent with prior observations, wild type Col-0 grown at 22 °C, showed increased hypocotyl elongation at 1 and 5 µM of PIC with 10 µM of PIC showing less (attenuated) induction (Gray et al., 1998, Schaefer et al., 2023) (**Fig. 5A & F**). For wild type at 28 °C, attenuation of growth began 5 µM PIC suggesting that either more auxin is natively produced, or plants show a higher auxin sensitivity (**Fig. 5A & G**). For *phyB-9* at 22° C, all PIC concentrations promoted hypocotyl elongation (**Fig. 5B, F**), whereas at 28° C, *phyB-9* hypocotyls showed attenuation at 10 µM PIC (**Fig. 5B & G**). As previously observed (Schaefer et al., 2023), *wdl4-3* seedlings at 22° C showed hyper-elongated at 1 and 5 µM PIC and attenuation at 10 µM PIC, similar to wild type (**Fig. 5C & F**). Interestingly, at 28° C, *wdl4-3* hyper-elongation was attenuated with 5 µM PIC, suggesting higher auxin sensitivity and/or production, and indicating a divergence form the *phyB-9* response (**Fig. 5C & G**). As previously reported (Franklin et al., 2011), *pif4* grown at 28° C was rescued back to wild type length by 1 µM PIC, (**Fig. 5G**, compare wild type + DMSO to *pif4* + 1µM PIC). Additionally, *pif4* showed an attenuated growth response at a higher PIC concentration at 28° C (**Fig. 5D, F, & G**). Elevated temperature in combination with 1 and 5 µM PIC restored hypocotyl elongation of the double *pif4 wdl4-3* seedlings to wild type (i.e., phenocopies *pif4*) length but was unable to promote hypocotyl growth to *wdl4-3* length (**Fig. 5E, F, & G**). These results indicate that auxin levels can account for PIF4-dependent hypocotyl elongation at 28° C, as previously demonstrated, but auxin does not account for the additional PIF4-dependent elongation observed in the absence of WDL4 function.

## Discussion

### WDL4 negatively regulates thermomorphogenesis and photomorphogenesis

Phytochromes play a fundamental role in plant biology, transducing both light and temperature signals to affect changes in growth and development (Krahmer and Fankhauser, 2024). PhyB has a central role in seedling de-etiolation with subsequent activities tied to photomorphogenesis and temperature acclimation (Quint et al., 2023, Romero-Montepaone et al., 2021). We observed that loss of WDL4 activity acts in every aspect of PhyB-dependent growth regulation, both cellular and developmental. When compared to the loss of function *phyb-9* mutant, the *wdl4-3* phenotypes appear specific and equivalent or more severe in nearly all respects. Moreover, ectopic WDL4-mNEON expression under a native promoter shows no gain of function phenotypes and rescues the *wdl4-3* mutant back to wildtype for growth and developmental phenotypes. These observations indicate that WDL4, a cytoplasmic microtubule associated protein, is required for PhyB-dependent regulation of both photomorphogenetic and thermomorphogenetic functions.

PhyB is synthesized in the cytoplasm in the Pr (red light absorbing) form and undergoes a reversible conformational change to the Pfr (far-red absorbing) after red light exposure (Reed et al., 1993, Chen et al., 2003, Burgie et al., 2021). Prior to translocation into the nucleus, PhyB can be post-translationally modified with effects on the efficiency of nuclear import and activity in the nucleus (Viczián and Nagy, 2024). Once imported, PhyB, in the Pfr form, interacts with PIFs, including PIF4, to repress transcription (Huq and Quail, 2002, Koini et al., 2009, Franklin et al., 2011). PIF repression is tied to the degradation of both PIFs and PhyB through a ubiquitin-mediated pathway (Leivar et al., 2012). Darkness, far-red light, or elevated ambient temperature will revert PhyB from the Pfr form back to the Pr form, leading to de-repression of PIFs and the transcription of hundreds of genes (Franklin et al., 2011, Sun et al., 2012, Oh et al., 2014, Xu and Zhu, 2021). WDL4 associates with cortical microtubules and has not been observed in the nucleus (Schaefer et al., 2023, Deng et al., 2021), suggesting that WDL4 acts on PhyB signaling either prior to PhyB nuclear translocation or, potentially, as a negative growth regulator after PIF4 activation. We propose from our observations that WDL4 acts on PhyB signaling prior to nuclear import to inhibit its negative regulation of PIF4. The proposal stems partly from observations that WDL4 rescues *wdl4-3* for red-light induced de-etiolation, a PhyB-dependent developmental switch involving substantial changes in gene expression (Leivar et al., 2008). Secondarily, we identified *wdl4-3* effects on negative gravitropism, thought to rely on cytoplasmic PhyB (Hu and Lagarias, 2024), and on bolting time, which requires PhyB interaction with PhyC (Sánchez-Lamas et al., 2016).

### WDL4 requires PIF4 and auxin signaling for function

PIF4 loss of function mutants show little morphological change in response to elevated temperatures (Franklin et al., 2011, Koini et al., 2009). Hypocotyls of *pif4* mutants show normal de-etiolation but remain short when grown in the light at elevated temperatures (Franklin et al., 2011, Koini et al., 2009). Exogenous auxin applied to *pif4* mutant seedlings at elevated temperature restores hypocotyl growth back to wild type TMG length, suggesting that auxin production downstream of PIF4 activation plays a foundational role in the temperature response pathway (Koini et al., 2009, Franklin et al., 2011). The TMG response is also blocked by the dominant *axr2-1* mutant, a nuclear localized auxin co-receptor that is not correctly degraded when auxin levels increase (Wilson et al., 1990, Schaefer et al., 2023). We found that double mutants with *wdl4-3* and either *axr2-1* or *pif4* failed to show the *wdl4-3* hyper-elongation phenotype at 22° or 28° C ((Schaefer et al., 2023); **Fig. 3 and Suppl. Fig. 3**). Moreover, application of auxin fails to elicit hypocotyl elongation in *wdl4-3 axr2-1* (Schaefer et al., 2023) but rescues growth back to wild type length in *pif4 wdl4-3* (**Fig. 5**). Together, these experiments provide evidence that auxin signaling is required for the correct hypocotyl TMG response and that the *wdl4-3* mutant is not interfering with the AXR2-dependent auxin response downstream of the PIF4 requirement in TMG.

Auxin plays a prominent role in many models for axial growth control owing to the capacity of plants to channel auxin flow and create gradients across tissues that correlate with cell expansion (Fendrych et al., 2016). WDL4 was proposed to control auxin trafficking, thus mediating auxin transport rather than production to modulate hypocotyl growth (Deng *et. al.,* 2021). Experiments using the auxin spatial transcriptional probe DR5_pro_:GFP failed to show an increase in auxin-dependent transcription in *wdl4-3* mutants and, consistent with the *pif4 wdl4-3* results, *wdl4-3* showed no shift in peak auxin sensitivity for axial growth (Schaefer et al., 2023). We observed that *phyb-9* hypocotyl growth exhibited a lower sensitivity to exogenous auxin concentration when compared to wild type and *wdl4-3*. However, neither *phyb-9* nor *wdl4-3* could be induced by exogenous auxin at 22° C to achieve the hypocotyl lengths observed at 28° C. Hence, auxin is clearly required for hypocotyl hyper-extension, but auxin level or trafficking does not account for the hyper-extended *phyb-9* or *wdl4-3* growth phenotypes at 28° C. Previous reports have shown that auxin-induced brassinosteriod (BR) signaling is important for temperature-induced hypocotyl elongation (Stavang et al., 2009, Delker et al., 2022). However, since *wdl4-3* seedlings have a similar response to BR as wild type (Schaefer et al 2023), we do not believe elevated BR signaling or sensitivity is responsible for temperature induced hyper-growth.

Why *phyb-9* and *wdl4-3* hypocotyls grow 2-4-fold longer than wild type in the light is not known. Crossing the *wdl4-3* mutation into the *pif4* background clearly blocked the *wdl4-3* hypocotyl elongation phenotype where multiple reports have shown that loss of PHYB in the *pif4* background results in hypocotyl hyper-extension (Huq and Quail, 2002, de Lucas et al., 2008). The *phyb-9* long hypocotyl phenotype is blocked when multiple PIFs are deleted, suggesting that PHYB normally represses PIF4 and an additional PIF family member(s) that can also elicit hyper-elongation. We interpret the *pif4 wdl4-3* double mutant phenotype to show that WDL4 likely acts specifically to facilitate PHYB interaction PIF4, where ectopic PIF4 expression is sufficient to induce hypocotyl hyper-elongation in light-grown plants (Sun et al., 2012).

The relationship of WDL4 to PIF4, and the positive effect of auxin downstream of PIF4, are not sufficient, however, to explain why loss of *wdl4-3* function results in very long hypocotyls. Recent work showing PhyB phosphorylation in the cytoplasm by plasma membrane localized FERONIA kinases suggests a possible connection to an auxin-independent growth regulation (Liu et al., 2023). FERONIA and related kinases retard growth by directly inactivating plasma membrane H^+^ ATPases through phosphorylation (Haruta et al., 2014). The additional phosphorylation of PhyB may provide a second route to modulating growth by changing PhyB activity levels or nuclear translocation (Chen et al., 2003, Liu et al., 2023, Viczián and Nagy, 2024). We note that the original *wdl4-3* hyper-elongation phenotype at 22° C, defined as continued growth from 4-7 dpg when wild type has paused (Schaefer et al., 2023), could align with a block to FERONIA activation, independent of auxin signaling. As of this publication, genetic interactions between PIF4 and FERONIA have not been determined under these conditions.

### A Microtubule Associated Protein in the PhyB Signaling Pathway

WDL4 belongs to a family of microtubule associated proteins that function in axial cell elongation (Yuen et al., 2003, Perrin et al., 2007, Liu et al., 2013, Sun et al., 2015, Deng et al., 2021, Okamoto et al., 2023, Schaefer et al., 2023). Loss of WDL4 activity results in hyper-elongation of the aerial organs, including hypocotyl and petioles (Deng et al 2021, Schaefer et al 2023), that have key roles in the adaptive thermomorphogenetic responses to elevated ambient temperature (Crawford et al., 2012). Axial growth also plays a major role in redirecting plant growth toward, or away from, environmental cues (Esmon et al., 2005, Vandenbussche et al., 2005, Verma et al., 2016). For example, preferencing axial growth to one side of a stem leads to bending in response to light signaling (Krahmer and Fankhauser, 2024). Controlling cell growth locally to correctly affect adaptive changes to plant shape requires significant coordination of physiological and molecular function. With this work, we show that WDL4 has a required role in connecting temperature and light signaling in the phytochrome pathway to plant morphogenesis both at seedling and adult life stages.

WDL4 localizes to cortical microtubules in the epidermal hypocotyl cells under both light and dark growth conditions (Schaefer et al., 2023). The absence of phenotype in dark-grown *wdl4-3* seedlings suggests that WDL4 is regulated to retard growth in the light, possibly through phosphorylations observed in proteomics studies (Vu et al., 2021, Arico et al., 2024). In turn, we proposed that WDL4 functions in the cytoplasm to promote PhyB suppression of PIF4. WDL4 shows physical association with syntaxins (Fujiwara et al., 2014, Deng et al., 2021) and therefore, WDL4 could help recruit or recycle proteins at the plasma membrane that modify PhyB prior to nuclear import related to PhyB stability or its ability to interact with other nuclear proteins. Alternatively, WDL4 could regulate the cytoplasmic interactions of PhyB with PIF4 to lower PIF4 levels or block nuclear entry, similar to proposals for PIF3 (Ni et al., 1999, Ni et al., 2013) and evidence for its cytoplasmic regulation (Hu and Lagarias, 2016, Hu and Lagarias, 2024).

## Methods

### Plant Growth Conditions

Seeds were surface sterilized (19:1 v/v 87% EtOH: 30% H_2_O_2_) and sown on ½ Murashige and Skoog (MS) medium supplemented with Gamborg’s Vitamins, 1% (w/v), plant specific agar (Sigma), pH 5.7 and without sucrose due to its effects on plant growth (Ohto et al., 2001). Seeds were stratified for 2-3 days in the dark at 4°C, moved to the appropriate light treatment to be grown horizontally.

For all high temperature seedling experiments, seedlings were grown in constant white light at 22°C for 3 days and then moved to 28°C for 3 additional days.

To test how elevated temperatures affect adult plant growth, seedlings were grown on plates for 10 days at 22°C in constant white light. Seedlings were then transferred to soil and grown at either 22°C or 28°C with a 12/12 hour light cycle. Each plant was followed and imaged every 7 days, up to 6 weeks using a Canon LSR. Side images of 3-week-old plants were used to measure petiole lengths and angle of hyponasty using ImageJ. The angle of hyponasty was measured setting the direction of growth as 0°, therefore a measurement closer to 0° represents a high amount of hyponasty and a measurement closer to 180° suggests non-hyponastic growth.

To determine seed germination rates, seeds were collected from individual plants after 6 weeks, and grown for 7 days at 22°C. At this time, any seedling that had broken the seed case was counted as successfully germinated. The total number of germinated seedlings was divided by the total number of seeds on the plate to determine the germination rate.

Supplemental far-red light was added using QBeam 2001 Solid State Lighting System (Quantum Devices, INC.) set to 20 µmol of 50/50 red/far-red light (670 nm/735 nm) in order to receive a reading on the light meter. Red wavelength light was then turned off. Seedlings were grown for 5 days before being analyzed. Each experimental round (5 total), genotype location on the plate was rotated (top, middle, bottom) to eliminate potential positional effects.

For red-light growth, after stratification seeds were exposed to white light at 22° C for 4-6 hours to promote germination. Seeds were then moved to a black box supplemented with horizontal 10 µmol of red light (670 nm) from the QBeam 2001 or wrapped in tinfoil for dark-grown controls. Seedlings were analyzed after 4 days. To reduce positional effects, genotype location on the plate was again moved during each experiment (7 total) to reduce potential positional effects.

All seedling plates were imaged with a Canon SLR camera. Hypocotyl, petiole, and root lengths and petiole angles were measured using ImageJ. Angle of hypocotyl growth was measured against the direction of gravity (range 0-180°, with 0° = vertical).

For Picloram (Hamaker et al., 1963) concentration series, seedlings were sown on nylon mesh (10 µM) laid on ½ MS and grown at 22° C in constant white light for 3 days. The nylon mesh with the seedlings was then transferred to the appropriate treatment plate (1/2 MS plus DMSO, 1, 5, or 10 µMol PIC) for 3 more days before analysis with ImageJ.

### Seed Lines

For all conditions, Columbia-0 (*Col-0*) was used as control/wild type. Other lines used are *wdl4-3* (SALK_015615, (Schaefer et al., 2023)), *phyB-9* (Reed et al., 1993), *pif4-101* (*pif4,* GARLIC 114-G06 (Huq and Quail, 2002)), 35s_pro_:WDL4-mNeon (Schaefer et al., 2023), and WDL4_pro:_WDL4-mNeon (Schaefer et al., 2023). To create the *pif4 wdl4-3* double mutant line, *pif4-101* was crossed into *wdl4-3* and verified by PCR analysis. (WDL4 WT F: GAAAGGTAAACCCGACAAAGG, WDL4 WT R: CCGAGATTCATGTCTCAGAGC, WDL4 tDNA F: GCGTGGACCGCTTGCTGCAACT) (*pif4* WT F: CTCGATTTCCGGTTATGG, *pif4* WT R: CAGACGGTTGATCATCTG; *pif4* tDNA F: GCATCTGAATTTCATAACCAATC)

### Statistical Analysis

Statistical analyses were performed at described in each figure legend.

### Accession Numbers

Sequence data from this article can be found in TAIR (arabidopsis.org) under the following accession numbers: WDL4 (AT2G35880), PIF4 (AT2G43010), PHYB (AT2G18790).

## Acknowledgements and Funding

We thank Roger Hangarter for numerous discussions and best methods for imaging adult plants. Funding for this work was provided by NSF-MCB1927504

## Author Contributions

K.S. planned, executed, and analyzed experiments and wrote the manuscript. S.L.S. contributed to the experimental design and analysis of the manuscript.

## Supplemental Data

**Supplemental Figure 1:**
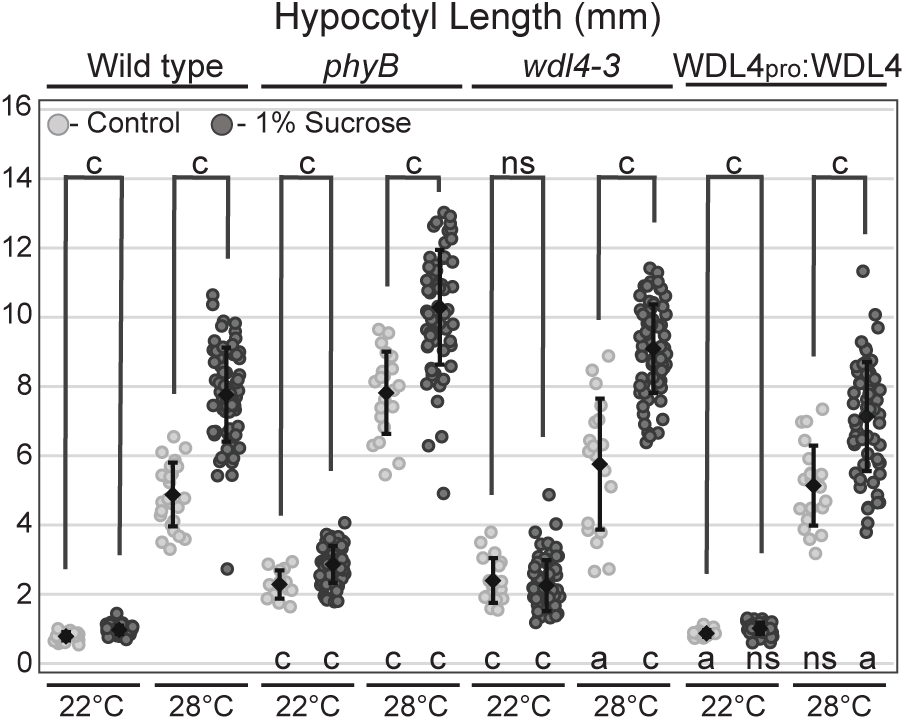
Sucrose enhances wild-type hyper-elongation at 28°C. Quantification of seedling hypocotyl lengths when grown on ½ MS plates without sucrose (light gray, n ≥ 14) or supplemented with 1% sucrose (dark gray, n ≥ 57). Seedlings were grown for 3 days at 22 °C then remained at 22 °C or shifted to 28 °C. Two-tailed student t-tests were used to compare measurements without or with 1% sucrose within a genotype (brackets and letters above) and between wild type and the mutant line with the same treatment (letters below). P-values: a < 0.05, b < 0.005, c < 0.0005, ns > 0.05. In dot plots, ♦ = mean value. Scale bars are standard deviation of the mean.

**Supplemental Figure 2:**
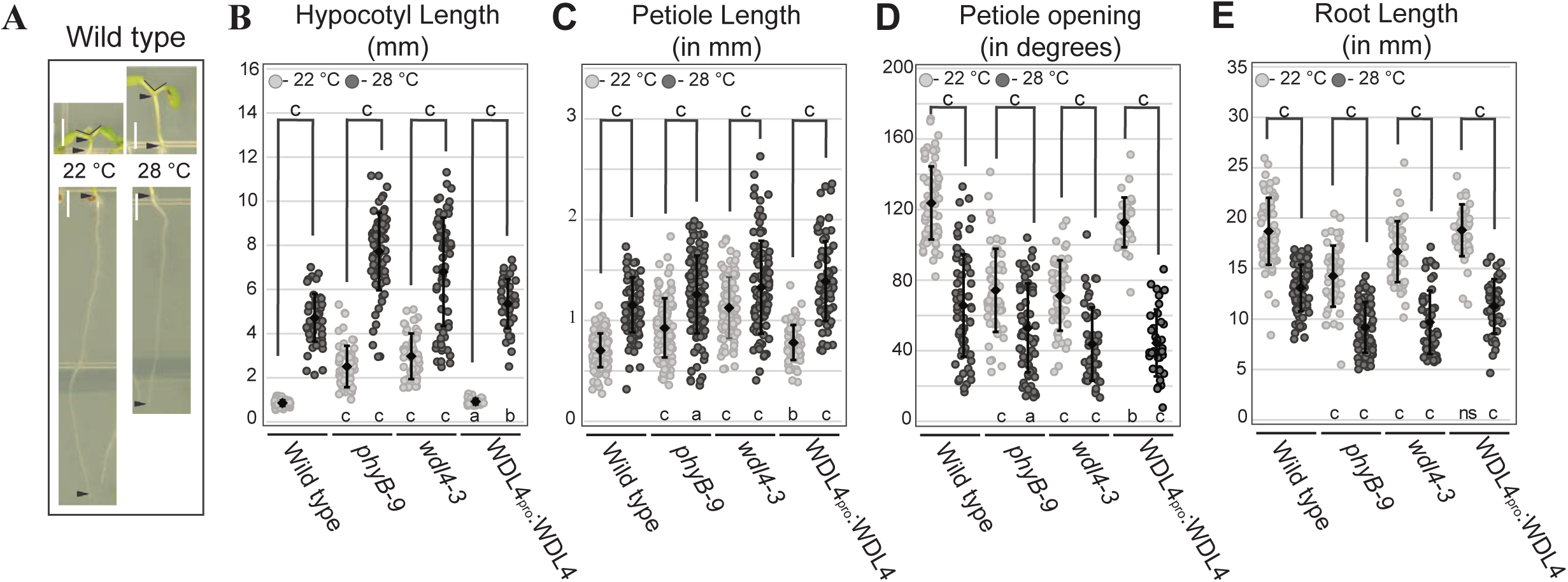
Thermomorphogenetic phenotype of *wdl4-3* seedlings. A) Representative images of 6 dpg wild type seedlings grown at 22 °C or 28 °C for days 4-6. Arrowheads show start and end of measurements for hypocotyl (top) or root (bottom) lengths. Large open V represents measurements for petiole opening. B) Hypocotyl lengths (n ≥ 39), C) Petiole lengths (n ≥ 75), D) angle of petiole opening (n ≥ 36), and E) root lengths (n ≥ 42) between seedlings grown at 22 °C or shifted to 28 °C at the start of day 4. WDL4pro:WDL4 = genomic WDL4 tagged with mNeon under the control of the WDL4 native promoter. Two-tailed student t-tests were used to compare measurements without or with 1% sucrose within a genotype (brackets and letters above) and between wild type and the mutant line with the same treatment (letters below). P-values: a < 0.05, b < 0.005, c < 0.0005, ns > 0.05. In dot plots, ♦ = mean value. Scale bars are standard deviation of the mean.

**Supplemental Figure 3:**
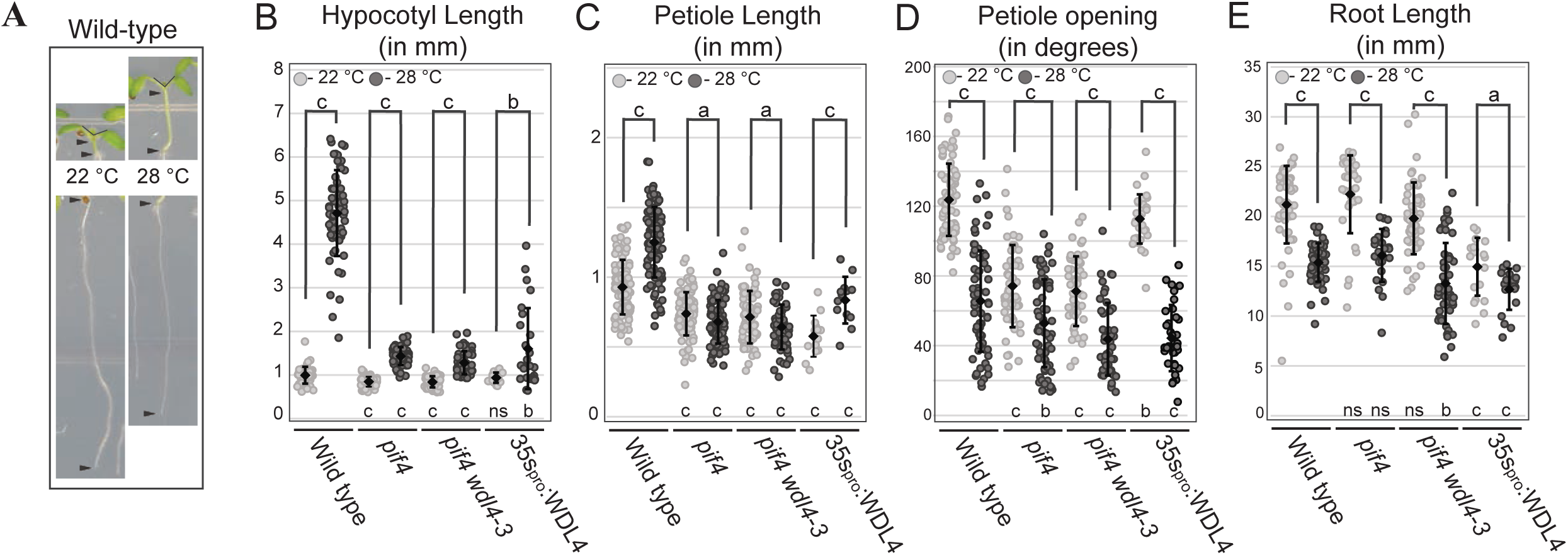
Thermomorphogenetic phenotype is diminished in *pif4 wdl4-3* seedlings. A) Representative images of 6 dpg wild type seedlings grown at 22 °C or 28 °C for days 4-6. Arrowheads show start and end of measurements for hypocotyl (top) or root (bottom) lengths. Large open V represents measurements for petiole opening. Quantification of B) Hypocotyl lengths (n ≥17 for 35spro:WDL4 and 53 for all others), C) Petiole lengths (n ≥14 for 35spro:WDL4 and 59 for all others), D) angle of petiole opening (n ≥ 22 for 35spro:WDL4 and 30 for all others), and E) root lengths (n ≥ 20 for 35spro:WDL4 and 25 for all others) between seedlings grown at 22 °C or shifted to 28 °C at the start of day 4. Two-tailed student t-tests were used to compare measurements without or with 1% sucrose within a genotype (brackets and letters above) and between wild type and the mutant line with the same treatment (letters below). P-values: a < 0.05, b < 0.005, c < 0.0005, ns > 0.05. In dot plots, ♦ = mean value. Scale bars are standard deviation of the mean. Note some scale bars are blocked by the mean value in B.

## References

Arico, D. S., Burachik, N. B., Wengier, D. L. & Mazzella, M. A. 2024. Arabidopsis hypocotyl growth in darkness requires the phosphorylation of a microtubule-associated protein. The Plant Journal, 118, 1815–1831.

Burgie, E. S., Gannam, Z. T. K., Mcloughlin, K. E., Sherman, C. D., Holehouse, A. S., Stankey, R. J. & Vierstra, R. D. 2021. Differing biophysical properties underpin the unique signaling potentials within the plant phytochrome photoreceptor families. Proceedings of the National Academy of Sciences, 118, e2105649118.

Casal, J. J. & Balasubramanian, S. 2019. Thermomorphogenesis. Annual Review of Plant Biology, 70, 321–346.

Chen, M., Schwab, R. & Chory, J. 2003. Characterization of the requirements for localization of phytochrome B to nuclear bodies. Proceedings of the National Academy of Sciences, 100, 14493–14498.

Crawford, A. J., Mclachlan, D. H., Hetherington, A. M. & Franklin, K. A. 2012. High temperature exposure increases plant cooling capacity. Current Biology, 22, R396–R397.

De Lucas, M., Davière, J.-M., RODRÍGUEZ-Falcón, M., Pontin, M., Iglesias-Pedraz, J. M., Lorrain, S., Fankhauser, C., Blázquez, M. A., Titarenko, E. & Prat, S. 2008. A molecular framework for light and gibberellin control of cell elongation. Nature, 451, 480–484.

Delker, C., Quint, M. & Wigge, P. A. 2022. Recent advances in understanding thermomorphogenesis signaling. Current Opinion in Plant Biology, 68, 102231.

Deng, J., Wang, X., Liu, Z. & Mao, T. 2021. The microtubule-associated protein WDL4 modulates auxin distribution to promote apical hook opening in Arabidopsis. Plant Cell, 33, 1927–1944.

Elliott, A. & Shaw, S. L. 2017. Microtubule Array Patterns Have a Common Underlying Architecture in Hypocotyl Cells. Plant Physiology, 176, 307–325.

Esmon, C. A., Pedmale, U. V. & Liscum, E. 2005. Plant tropisms: providing the power of movement to a sessile organism. Int J Dev Biol, 49, 665–74.

Fendrych, M., Leung, J. & Friml, J. 2016. TIR1/AFB-Aux/IAA auxin perception mediates rapid cell wall acidification and growth of Arabidopsis hypocotyls. eLife, 5, e19048.

Franklin, K. A., Lee, S. H., Patel, D., Kumar, S. V., Spartz, A. K., Gu, C., Ye, S., Yu, P., Breen, G., Cohen, J. D., Wigge, P. A. & Gray, W. M. 2011. PHYTOCHROME-INTERACTING FACTOR 4 (PIF4) regulates auxin biosynthesis at high temperature. Proceedings of the National Academy of Sciences, 108, 20231–20235.

Franklin, K. A. & Whitelam, G. C. 2005. Phytochromes and Shade-avoidance Responses in Plants. Annals of Botany, 96, 169–175.

Fujiwara, M., Uemura, T., Ebine, K., Nishimori, Y., Ueda, T., Nakano, A., Sato, M. H. & Fukao, Y. 2014. Interactomics of Qa-SNARE in Arabidopsis thaliana. Plant Cell Physiol, 55, 781–9.

Gendreau, E., Traas, J., Desnos, T., Grandjean, O., Caboche, M. & Höfte, H. 1997. Cellular Basis of Hypocotyl Growth in Arabidopsis thaliana. Plant Physiology, 114, 295–305.

Gray, W. M., Östin, A., Sandberg, G., Romano, C. P. & Estelle, M. 1998. High temperature promotes auxin-mediated hypocotyl elongation in *Arabidopsis*. Proceedings of the National Academy of Sciences, 95, 7197–7202.

Hamaker, J. W., Johnston, H., Martin, R. T. & Redemann, C. T. 1963. A Picolinic Acid Derivative: A Plant Growth Regulator. Science, 141, 363.

Haruta, M., Sabat, G., Stecker, K., Minkoff, B. B. & Sussman, M. R. 2014. A Peptide Hormone and Its Receptor Protein Kinase Regulate Plant Cell Expansion. Science, 343, 408–411.

Hatfield, J. L. & Prueger, J. H. 2015. Temperature extremes: Effect on plant growth and development. Weather and Climate Extremes, 10, 4–10.

Hu, W. & Lagarias, J. C. 2016. A Tightly Regulated Genetic Selection System with Signaling-Active Alleles of Phytochrome B. Plant Physiology, 173, 366–375.

Hu, W. & Lagarias, J. C. 2024. A cytosol-tethered YHB variant of phytochrome B retains photomorphogenic signaling activity. Plant Molecular Biology, 114, 72.

Hughes, J. 2013. Phytochrome Cytoplasmic Signaling. Annual Review of Plant Biology, 64, 377–402.

Huq, E. & Quail, P. H. 2002. Pif4, a phytochrome&#x2010;interacting bHLH factor, functions as a negative regulator of phytochrome B signaling in *Arabidopsis*. The EMBO Journal, 21, 2441–2450.

Jung, J.-H., Domijan, M., Klose, C., Biswas, S., Ezer, D., Gao, M., Khattak, A. K., Box, M. S., Charoensawan, V., Cortijo, S., Kumar, M., Grant, A., Locke, J. C. W., Schäfer, E., Jaeger, K. E. & Wigge, P. A. 2016. Phytochromes function as thermosensors in *Arabidopsis*. Science, 354, 886–889.

Kim, C., Kwon, Y., Jeong, J., Kang, M., Lee, G. S., Moon, J. H., Lee, H.-J., Park, Y.-I. & Choi, G. 2023. Phytochrome B photobodies are comprised of phytochrome B and its primary and secondary interacting proteins. Nature Communications, 14, 1708.

Koini, M. A., Alvey, L., Allen, T., Tilley, C. A., Harberd, N. P., Whitelam, G. C. & Franklin, K. A. 2009. High Temperature-Mediated Adaptations in Plant Architecture Require the bHLH Transcription Factor PIF4. Current Biology, 19, 408–413.

Krahmer, J. & Fankhauser, C. 2024. Environmental Control of Hypocotyl Elongation. Annual Review of Plant Biology, 75, null.

Lee, S., Zhu, L. & Huq, E. 2021. An autoregulatory negative feedback loop controls thermomorphogenesis in Arabidopsis. PLOS Genetics, 17, e1009595.

Legris, M., Klose, C., Burgie, E. S., Rojas, C. C. R., Neme, M., Hiltbrunner, A., Wigge, P. A., Schäfer, E., Vierstra, R. D. & Casal, J. J. 2016. Phytochrome B integrates light and temperature signals in *Arabidopsis*. Science, 354, 897–900.

Leivar, P., Monte, E., Al-Sady, B., Carle, C., Storer, A., Alonso, J. M., Ecker, J. R. & Quail, P. H. 2008. The Arabidopsis Phytochrome-Interacting Factor Pif7, Together with PIF3 and Pif4, Regulates Responses to Prolonged Red Light by Modulating phyB Levels. The Plant Cell, 20, 337–352.

Leivar, P., Monte, E., Cohn, M. M. & Quail, P. H. 2012. Phytochrome Signaling in Green Arabidopsis Seedlings: Impact Assessment of a Mutually Negative phyB–PIF Feedback Loop. Molecular Plant, 5, 734–749.

Liu, X., Jiang, W., Li, Y., Nie, H., Cui, L., Li, R., Tan, L., Peng, L., Li, C., Luo, J., Li, M., Wang, H., Yang, J., Zhou, B., Wang, P., Liu, H., Zhu, J.-K. & Zhao, C. 2023. FERONIA coordinates plant growth and salt tolerance via the phosphorylation of phyB. Nature Plants, 9, 645–660.

Liu, X., Qin, T., Ma, Q., Sun, J., Liu, Z., Yuan, M. & Mao, T. 2013. Light-regulated hypocotyl elongation involves proteasome-dependent degradation of the microtubule regulatory protein WDL3 in Arabidopsis. Plant Cell, 25, 1740–55.

Lorrain, S., Allen, T., Duek, P. D., Whitelam, G. C. & Fankhauser, C. 2008. Phytochrome-mediated inhibition of shade avoidance involves degradation of growth-promoting bHLH transcription factors. The Plant Journal, 53, 312–323.

Lu, X.-D., Zhou, C.-M., Xu, P.-B., Luo, Q., Lian, H.-L. & Yang, H.-Q. 2015. Red-Light-Dependent Interaction of phyB with SPA1 Promotes COP1–SPA1 Dissociation and Photomorphogenic Development in Arabidopsis. Molecular Plant, 8, 467–478.

Ni, M., Tepperman, J. M. & Quail, P. H. 1999. Binding of phytochrome B to its nuclear signalling partner PIF3 is reversibly induced by light. Nature, 400, 781–784.

Ni, W., Xu, S.-L., Chalkley, R. J., Pham, T. N. D., Guan, S., Maltby, D. A., Burlingame, A. L., Wang, Z.-Y. & Quail, P. H. 2013. Multisite Light-Induced Phosphorylation of the Transcription Factor PIF3 Is Necessary for Both Its Rapid Degradation and Concomitant Negative Feedback Modulation of Photoreceptor phyB Levels in Arabidopsis The Plant Cell, 25, 2679–2698.

Oh, E., Zhu, J.-Y., Bai, M.-Y., Arenhart, R. A., Sun, Y. & Wang, Z.-Y. 2014. Cell elongation is regulated through a central circuit of interacting transcription factors in the Arabidopsis hypocotyl. eLife, 3, e03031.

Ohto, M.-A., Onai, K., Furukawa, Y., Aoki, E., Araki, T. & Nakamura, K. 2001. Effects of Sugar on Vegetative Development and Floral Transition in Arabidopsis. Plant Physiology, 127, 252–261.

Okamoto, T., Motose, H. & Takahashi, T. 2023. Microtubule-associated proteins WDL5 and WDL6 play a critical role in pollen tube growth in Arabidopsis thaliana. Plant Signaling & Behavior, 18, 2281159.

Perrin, R. M., Wang, Y., Yuen, C. Y. L., Will, J. & Masson, P. H. 2007. WVD2 is a novel microtubule-associated protein in Arabidopsis thaliana. The Plant Journal, 49, 961–971.

Pucciariello, O., Legris, M., COSTIGLIOLO Rojas, C., Iglesias, M. J., Hernando, C. E., Dezar, C., Vazquez, M., Yanovsky, M. J., Finlayson, S. A., Prat, S. & Casal, J. J. 2018. Rewiring of auxin signaling under persistent shade. Proceedings of the National Academy of Sciences, 115, 5612–5617.

Quint, M., Delker, C., Balasubramanian, S., Balcerowicz, M., Casal, J. J., Castroverde, C. D. M., Chen, M., Chen, X., DE Smet, I., Fankhauser, C., Franklin, K. A., Halliday, K. J., Hayes, S., Jiang, D., Jung, J.-H., Kaiserli, E., Kumar, S. V., Maag, D., Oh, E., Park, C.-M., Penfield, S., Perrella, G., Prat, S., Reis, R. S., Wigge, P. A., Willige, B. C. & Van Zanten, M. 2023. 25 Years of thermomorphogenesis research: milestones and perspectives. Trends in Plant Science, 28, 1098–1100.

Quint, M., Delker, C., Franklin, K. A., Wigge, P. A., Halliday, K. J. & VAN Zanten, M. 2016. Molecular and genetic control of plant thermomorphogenesis. Nature Plants, 2, 15190.

Reed, J. W., Nagatani, A., Elich, T. D., Fagan, M. & Chory, J. 1994. Phytochrome A and Phytochrome B Have Overlapping but Distinct Functions in Arabidopsis Development. Plant Physiology, 104, 1139–1149.

Reed, J. W., Nagpal, P., Poole, D. S., Furuya, M. & Chory, J. 1993. Mutations in the gene for the red/far-red light receptor phytochrome B alter cell elongation and physiological responses throughout Arabidopsis development. The Plant Cell, 5, 147–157.

Romero-Montepaone, S., Sellaro, R., Esteban Hernando, C., Costigliolo-Rojas, C., Bianchimano, L., Ploschuk, E. L., Yanovsky, M. J. & Casal, J. J. 2021. Functional convergence of growth responses to shade and warmth in Arabidopsis. New Phytologist, 231, 1890–1905.

Sánchez-Lamas, M., Lorenzo, C. D. & Cerdán, P. D. 2016. Bottom-up Assembly of the Phytochrome Network. PLOS Genetics, 12, e1006413.

Schaefer, K., Cairo Baza, A., Huang, T., Cioffi, T., Elliott, A. & Shaw, S. L. 2023. WAVE-DAMPENED2-LIKE4 modulates the hyper-elongation of light-grown hypocotyl cells. Plant Physiology, 192, 2687–2702.

Spartz, A. K., Lee, S. H., Wenger, J. P., Gonzalez, N., Itoh, H., Inzé, D., Peer, W. A., Murphy, A. S., Overvoorde, P. J. & Gray, W. M. 2012. The SAUR19 subfamily of SMALL AUXIN UP RNA genes promote cell expansion. The Plant Journal, 70, 978–990.

Stavang, J. A., Gallego-Bartolomé, J., Gómez, M. D., Yoshida, S., Asami, T., Olsen, J. E., GARCÍA-Martínez, J. L., Alabadí, D. & Blázquez, M. A. 2009. Hormonal regulation of temperature-induced growth in Arabidopsis. The Plant Journal, 60, 589–601.

Sun, J., Ma, Q. & Mao, T. 2015. Ethylene Regulates the Arabidopsis Microtubule-Associated Protein WAVE-DAMPENED2-LIKE5 in Etiolated Hypocotyl Elongation. Plant Physiol, 169, 325–37.

Sun, J., Qi, L., Li, Y., Chu, J. & Li, C. 2012. PIF4–Mediated Activation of YUCCA8 Expression Integrates Temperature into the Auxin Pathway in Regulating Arabidopsis Hypocotyl Growth. PLOS Genetics, 8, e1002594.

True, J. H. & Shaw, S. L. 2020. Exogenous Auxin Induces Transverse Microtubule Arrays Through TRANSPORT INHIBITOR RESPONSE1/AUXIN SIGNALING F-BOX Receptors. Plant Physiol, 182, 892–907.

Ulmasov, T., Hagen, G. & Guilfoyle, T. J. 1999. Activation and repression of transcription by auxin-response factors. Proceedings of the National Academy of Sciences, 96, 5844–5849.

Van Zanten, M., Voesenek, L. A. C. J., Peeters, A. J. M. & Millenaar, F. F. 2009. Hormone- and Light-Mediated Regulation of Heat-Induced Differential Petiole Growth in Arabidopsis. Plant Physiology, 151, 1446–1458.

Vandenbussche, F., Verbelen, J.-P. & VAN DER Straeten, D. 2005. Of light and length: Regulation of hypocotyl growth in Arabidopsis. BioEssays, 27, 275–284.

Verma, V., Ravindran, P. & Kumar, P. P. 2016. Plant hormone-mediated regulation of stress responses. BMC Plant Biology, 16, 86.

Viczián, A. & Nagy, F. 2024. Phytochrome B phosphorylation expanded: site-specific kinases are identified. New Phytologist, 241, 65–72.

Vu, L. D., Xu, X., Zhu, T., Pan, L., Van Zanten, M., De Jong, D., Wang, Y., Vanremoortele, T., Locke, A. M., Van De Cotte, B., De Winne, N., Stes, E., Russinova, E., De Jaeger, G., Van Damme, D., Uauy, C., Gevaert, K. & De Smet, I. 2021. The membrane-localized protein kinase MAP4K4/TOT3 regulates thermomorphogenesis. Nature Communications, 12, 2842.

Wang, Z.-Y., Bai, M.-Y., Oh, E. & Zhu, J.-Y. 2012. Brassinosteroid Signaling Network and Regulation of Photomorphogenesis. Annual Review of Genetics, 46, 701–724.

Wilson, A. K., Pickett, F. B., Turner, J. C. & Estelle, M. 1990. A dominant mutation in Arabidopsis confers resistance to auxin, ethylene and abscisic acid. Mol Gen Genet, 222, 377–83.

Xu, Y. & Zhu, Z. 2021. PIF4 and PIF4-Interacting Proteins: At the Nexus of Plant Light, Temperature and Hormone Signal Integrations. International Journal of Molecular Sciences, 22, 10304.

Yuen, C. Y., Pearlman, R. S., Silo-Suh, L., Hilson, P., Carroll, K. L. & Masson, P. H. 2003. WVD2 and WDL1 modulate helical organ growth and anisotropic cell expansion in Arabidopsis. Plant Physiol, 131, 493–506.

